# The Neural Basis of Shared Preference Learning

**DOI:** 10.1101/570762

**Authors:** Harry Farmer, Uri Hertz, Antonia Hamilton

## Abstract

During our daily lives, we often learn about the similarity of the traits and preferences of others to our own and use that information during our social interactions. However, it is unclear how the brain represents similarity between the self and others. One possible mechanism is to track similarity to oneself regardless of the identity of the other (Similarity account); an alternative is to track each confederate in terms of consistency of the similarity to the self, with respect to the choices they have made before (consistency account). Our study combined fMRI and computational modelling of reinforcement learning (RL) to investigate the neural processes that underlie learning about preference similarity. Participants chose which of two pieces of artwork they preferred and saw the choices of one confederate who usually shared their preference and another who usually did not. We modelled neural activation with RL models based on the similarity and consistency accounts. Data showed more brain regions whose activity pattern fits with the consistency account, specifically, areas linked to reward and social cognition. Our findings suggest that impressions of other people can be calculated in a person-specific manner which assumes that each individual behaves consistently with their past choices.

## 1. Introduction

The ability to rapidly form and update our impressions about other people is a vital skill in navigating our complex social world. During our daily lives, we frequently learn about the traits and preferences of other people and use that information to inform our social interactions. For example, if I like Monet, and walk around a gallery with Alice who comments favourably on the Monet, while Beth prefers the Gauguin, I will learn that Alice’s preferences are similar to mine and may like Alice more or feel closer to her. However, the neural mechanisms which govern our learning of the relationship between our preferences and those of others are currently unclear.

Theoretically, there are at least two ways that people might learn about others. They might track each person as a unique individual, inferring their behaviour and predicting future choices based only on their previous actions. This has commonly been studied within the context of impression formation (Hackel, Doll, & Amodio, 2015; Mende-Siedlecki, 2018). Alternatively, people might track others in terms of their similarity or difference to the self, as predicted by theories of anchoring and adjustment (D. R. Ames, 2004; Tamir & Mitchell, 2012). In the current study, we combined fMRI and computational modelling of reinforcement learning (RL) to investigate the neural processes that underlie learning about the similarity of others’ preferences to our own.

Researchers investigating impression formation have sought to determine which brain areas respond when we learn about other people and when our expectations of others are violated. Most have done this by providing participants with some information about a novel person and then presenting either consistent information which simply confirms the previous impression or inconsistent which requires participants to update their impressions. These studies have shown increased activity in regions like the precuneus/posterior cingulate cortex (PCC), the temporal-parietal junction (TPJ) and the dorsomedial prefrontal cortex (dmPFC) when receiving inconsistent vs. consistent information about another person’s moral behaviour (Hughes, Zaki, & Ambady, 2017; Mende-Siedlecki, Baron, & Todorov, 2013; Mende-Siedlecki & Todorov, 2016), competence (D. L. Ames & Fiske, 2013; Bhanji & Beer, 2013), traits (Hackel et al., 2015; Ma et al., 2012; Van der Cruyssen, Heleven, Ma, Vandekerckhove, & Van Overwalle, 2015), and political beliefs (Cloutier, Gabrieli, Young, & Ambady, 2011). These regions are key nodes in the “mentalising” network which is activated when thinking about the beliefs, preferences and intentions of others (Adolphs, 2009; Frith & Frith, 2012; Schilbach, 2015; Van Overwalle, 2009).

The increased activation to inconsistent information seen in the mentalising network is reminiscent of the prediction error signal seen in RL models. These signals compute the expectation of a future outcome (or reward) as being a function of the current expectation plus the product of the learning rate and the prediction error, i.e. the difference between the last expected and actual outcome (Behrens, Hunt, & Rushworth, 2009; Ruff & Fehr, 2014). This has led researchers to suggest that these regions may be involved in calculating social prediction errors (Hertz et al., 2017; Mende-Siedlecki, Cai, & Todorov, 2013). Previous studies have investigated this possibility directly, using computational modelling to parametrically track prediction error from trial to trial and have found evidence of social prediction error tracking in the dmPFC, the TJP, the STS, the medial temporal gyrus (MTG), ventrolateral PFC (vlPFC) and the precuneus (Behrens, Hunt, Woolrich, & Rushworth, 2008; Hackel et al., 2015; Stanley, 2016). These and other findings have led some researchers (e.g. Bach & Schenke, 2017; Joiner, Piva, Turrin, & Chang, 2017) to argue that predictive processing plays a key role in social cognition.

However, most of the studies examining social prediction errors have considered cases where participants learn about a single individual, and do not manipulate the relationship between that individual and the self. In the real world, we are able to learn about many different individuals and we often make judgements about others in relation to ourselves. A distinct literature has examined the role of self-similarity in impression formation (Boer et al., 2011; Montoya & Horton, 2013). A particular focus is the idea that self-similarity can lead to liking and affiliation. Numerous studies have shown that those we perceive as similar to us in terms of traits (Paunonen & Hong, 2013), attitudes (Montoya & Horton, 2013) and preferences (Boer et al., 2011) tend to be evaluated more favourably than those perceived as different. Several studies have presented evidence for a ventral-dorsal gradient in the mPFC when processing similar and dissimilar others with greater activation for those similar to the self in the vmPFC and greater processing of dissimilar others in the dmPFC (Denny, Kober, Wager, & Ochsner, 2012; Sul et al., 2015). However, few studies have examined evaluation of self and other in terms of social learning.

Tracking the relationship between our own preferences and those of others is also important in social influence, in which we change our preferences to be more in line with those of the other people (Moutoussis, Dolan, & Dayan, 2016; Narayan, Rao, & Saunders, 2011). Several studies have shown that key nodes of the mentalising network including the mPFC (Garvert, Moutoussis, Kurth-Nelson, Behrens, & Dolan, 2015; Nook & Zaki, 2015; Zaki, Schirmer, & Mitchell, 2011), the precuneus and the TPJ (Berns, Capra, Moore, & Noussair, 2010; Campbell-Meiklejohn, Bach, Roepstorff, Dolan, & Frith, 2010; Welborn et al., 2016) play a key role in social influence.

The current study aims to test how the brain tracks and learns about other people from the self-similarity of their choices, which will help with understanding the link between social prediction errors and theories of the sense of self and the ability to build affiliations. To do this we adapted reinforcement learning models to investigate the tracking of similarity of choice between self and others. It is important to note here that we are not claiming that the tracking of similarity is necessarily linked to reward based reinforcement in a direct manner. Rather, we are interested in the use of RL models due to their utility in modelling the accumulation of information and evidence over time. This allows us to look at how the brain represents confirming and disconfirming information about others similarity to ourselves. For a similar approach applied to the learning of others’ traits see Zaki, Kallman, Wimmer, Ochsner and Shohamy (2016).

Our task created a context in which participants chose which painting they prefer (an arbitrary aesthetic choice) and then learn the preferences of two other confederates for the same paintings. Using fMRI and computational modelling, we can identify which brain systems track others’ preferences relative to self-preferences in a trial-by-trial manner, and whether the two confederates are both tracked in respect to their similarity to self or in respect to their previous behaviour. In each trial our participants saw two paintings and indicated which they preferred (Figure 1A). They then saw the preferences of two confederates, a similar confederate (Sim) who chose the same painting 75% of the time and a different confederate (Diff) who chose the same painting only 25% of the time. We then build RL models that give us the prior probability of the confederates’ choice and the prediction error of their actual choice separately for each trial and each confederate, based on the trial sequence experienced by the participants, and use this to localise brain regions where the BOLD signal tracks the model parameters.

**Figure 1.**
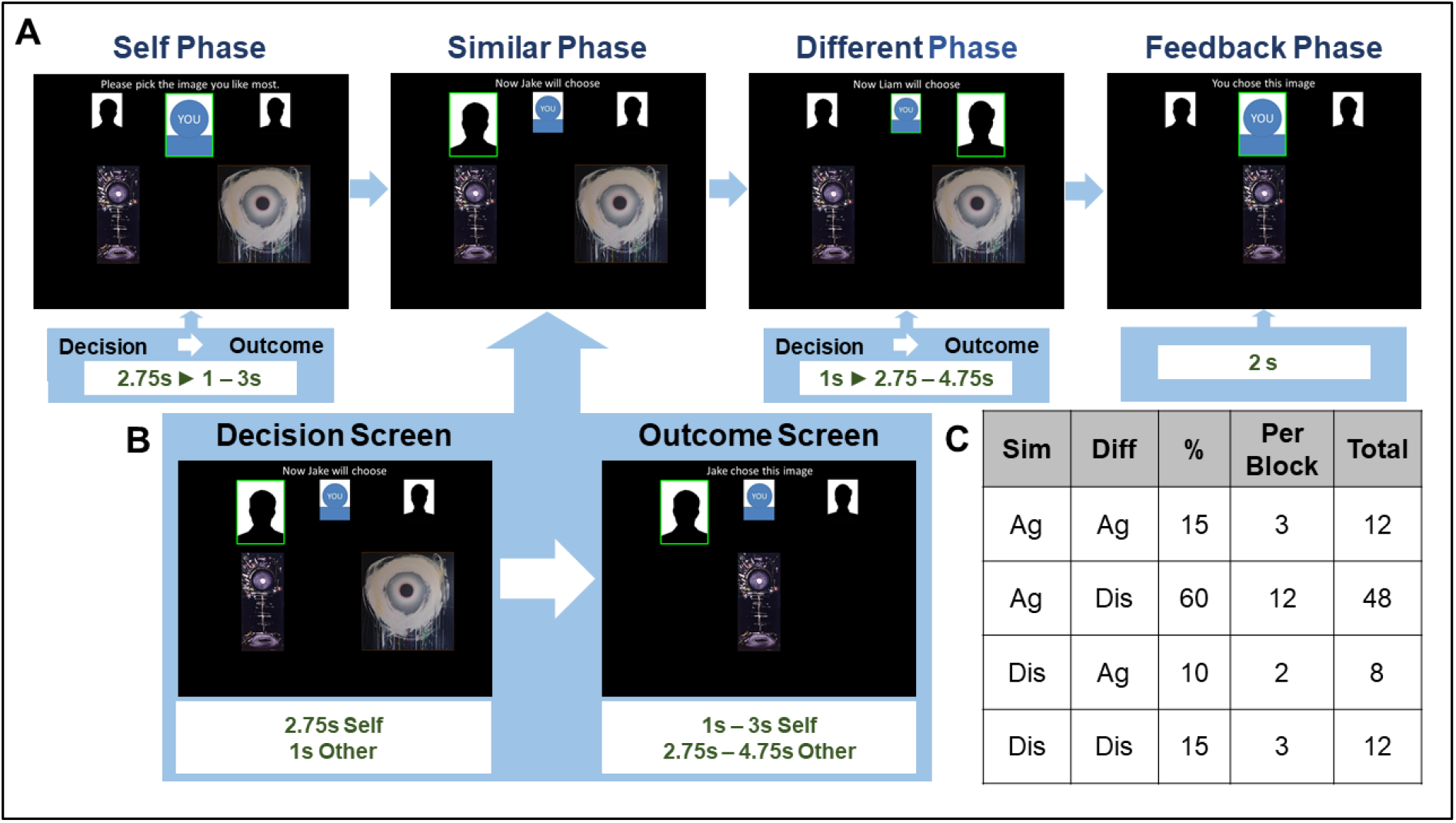
Outline of experimental trial structure and number of trials per condition. A. Each trial has four phases (self, similar, different, feedback). Each of the first three phases is composed of a decision screen and an outcome screen. B) An expanded view of the decision and outcome screens is illustrated for the Similar phase in but the same structure was used for the Self and Different phases. During the decision screen participants either chose their own preferred painting (Self phase) or waited to see the choice of the confederate (Similar & Different phases). During the outcome screen participants either saw the painting they had chosen (Self phase) or the painting the confederate chose (Similar & Different phases). During the feedback phase participants saw their own choice again. Durations for each screen are shown in green, s = seconds. The order of similar and different confederates was counterbalanced across trials. C) This table shows the breakdown of the four possible combinations of choices by percentage, block and total number of trials. Sim = similar, Diff = different, Ag = agree with participant s choice, Dis = disagree with participants choice. Note in the actual study we used images from the Karolinska Directed Emotional Faces database for the confederates.

When considering how the brain might track accumulated similarity and prediction error for the two confederates we modelled two possible approaches (see Figure 2A), which we term the ‘Similarity approach and the ‘Consistency approach. In the Similarity approach, each confederate is tracked in relation to one’s own preferences, on a single dimension of ‘distance from me’. We operationalised this approach in a RL model which tracks each confederates’ similarity to the participant’s preferences over all trials, which we term Accumulated Similarity (AS). In this model each trial provides a new observation of similarity and thus a signed similarity prediction error (PE_Sim), where positive values indicating greater-than-expected similarity and negative values indicating greater-than-expected dissimilarity.

**Figure 2.**
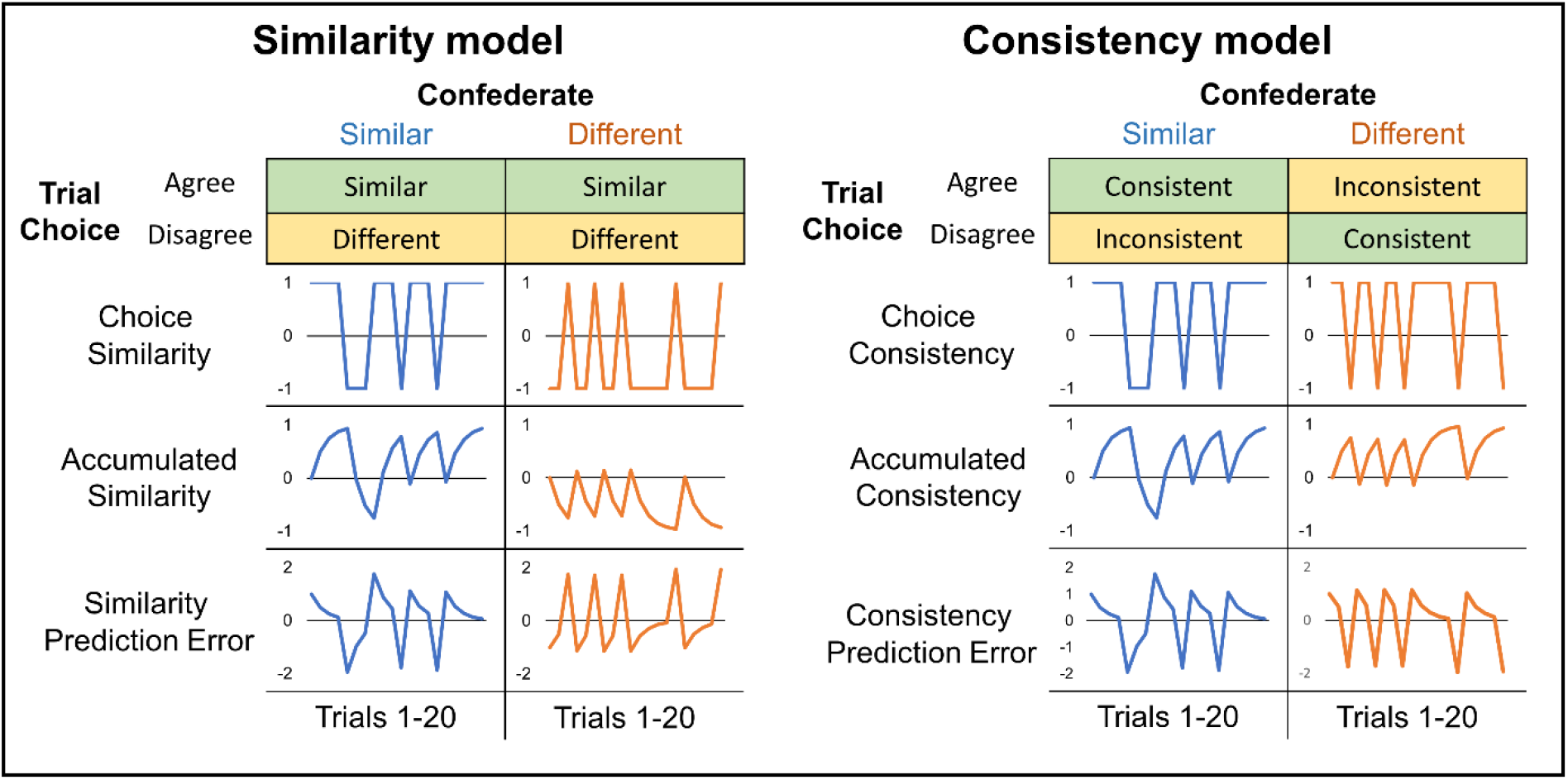
Two possible ways that the choices of people with similar and different overall preferences may be tracked in the brain. Diagram showing similarity and consistency models. For each model we show the relationship between confederate trait (similar/different) and choice on each trial relative to the participant (agree/disagree) combines to define whether that trail is coded as similar/consistent. Below that we display how choice input, accumulated similarity/consistency and prediction error for the similar and different confederates change across one session (20 trials).

In the Consistency approach, we assume that each confederate is tracked with respect to their own past track record of behaviour, which here means their track record of agreeing with the participant or of disagreeing. This was operationalised in a RL model which tracks each confederates’ consistency to their overall tendency to make similar or different choices to that of the participant over all trials, which we term Accumulated Consistency (AC). In this model the prediction errors are therefore not be in relation to that choice’s similarity to the self (similar/dissimilar) but to the confederate’s overall similarity to the self, inferred from previous pattern of choices (consistent/inconsistent) thus consistency prediction errors (PE_Con). In this case negative PE_Con will represent greater than expected inconsistent choices, regardless of the similarity to the participants’ own choice.

Importantly, these two models predict a completely different pattern of brain activity in our experimental design, as similar confederate’s consistent responses and different confederate consistent responses can have the same sign (consistency model) or opposite sign (similarity model) (see Figure 2).

## 2 Methods

### 2.1 Design

In our study participants tracked the choices of two confederates on multiple trials, in relation to their own choices. On each trial, the participant and two confederates indicated which of two paintings they preferred. The similar confederate (Sim) chose the same painting as the participant in 75% of all trials while the different confederate (Diff) only chose the same painting in 25% of trials.

### 2.2 Participants

Twenty-five participants (mean age ± *SD:* 25.1 ± 5.7, 11 male) took part in this study which was approved by The University College London, Institute of Cognitive Neuroscience Research Department’s Ethics Committee. All participants gave their informed consent to participate and were paid for their participation. All participants were right-handed and were screened for neurological disorders. Due to technical issues, behavioural data was lost for 7 participants. Therefore, our final sample size for behavioural analysis was n=18. As we did not use behavioural data for model fitting, and choice’s similarity was tracked regardless of the participants’ choices, this issue did not impact on the fMRI analysis so the full sample n=25 was used for fMRI analysis.

### 2.3 Procedure

#### 2.3.1 Experimental Task

The main task in this study was an aesthetic choice task. Participants were told that in each trial they would see a pair of paintings (see Supplementary Materials for details of paintigs’ pretesting & matching) and would have to choose which painting they preferred. They were told that other participants had previously indicated which of the paintings they preferred and that they would see those participant’s choices during the study.

Prior to entering the scanner participants completed a training block of the task consisting of 10 trials, to ensure that they understood the task. In the training block, they saw two confederates of the opposite gender to themselves, and choices were made in the order: Confederate 1, Participant, Confederate 2. This was to create the belief that the order of making choices was random and that each confederate made choices independently. In fact, in the experimental trials, the choices of the confederates were determined from the participant choices to create appropriate levels of similarity. After the training, participants learnt the names of the confederates with whom they would do the experimental task and rated their faces for similarity, likeability and attractiveness, using a 10-point scale. In the experimental blocks, participants saw these two confederates of the same gender as them and the participants’ choice always came before the choices of the two confederates. Images of the confederates were taken from the Karolinska Directed Emotional Face database (Lundqvist, Flykt, and Öhman 1998) but participants were informed that these photos were stand in images for the actual other participants.

#### 2.3.2 Trial Structure

Each trial was divided into four phases (see Figure 2A). The first three phases were split into two screens, a *decision screen* and an *outcome screen* (see Figure 2B). In the Self-phase participants were shown a pair of paintings on the *decision* screen and had 2.75 seconds to choose which they preferred using the left and right buttons on a response box. They then saw an *outcome* screen displaying their preferred painting for a jittered interval (1-3 seconds). In the Similar-phase, participants first saw a 1-second *decision* screen which displayed the pair of paintings along with an indicator of which confederate was choosing. This was followed by an *outcome* screen which displayed the confederate’s preferred painting for a jittered interval (2.75 −4.75 seconds). In the Different-phase, participants again saw a *decision* screen with an indicator that the other confederate was choosing, followed by a jittered *outcome* screen displaying that confederate’s preferred painting. The order of the similar & different phases was pseudorandomised by trial. Finally, each trial contained a *feedback* phase in which participants again saw their own choice for an interval of 2-seconds.

Participants completed 4 sessions of 20 trials (see Figure 1C for a breakdown of trial types by block), and at the end of each block, they rated the similarity, likeability and attractiveness of each confederate using a 10-point scale. Each block was one scanner session with a short break between.

Varying the intervals of the outcome screen in the three choice phases achieved an effective temporal sampling resolution much finer than one TR for each of these periods. The lengths of the intervals were uniformly distributed for each period, ensuring that Evoked Haemodynamic Responses time-locked to the events were sampled evenly across the time period following each choice period.

### 2.4 Image Acquisition and Data Analysis

A 1.5 T Siemens TIM Avanto scanner with a 32-channel head coil was used to acquire both T1-weighted structural images and T2*-weighted echoplanar images using the multiband method (64×64 pixels; 3.2×3.2 mm; echo time, 55 ms, multiband factor=2) with blood oxygen level-dependent (BOLD) contrast. Each volume comprised 40 axial slices (3.2 mm thick, oriented approximately to the anterior commissure–posterior commissure plane), covering most of the brain but omitting inferior portions of the cerebellum. Functional scans were acquired in four sessions, each comprising 222 volumes (~7.4 min). Volumes were acquired continuously with an effective repetition time of 2s per volume. The first four volumes in each session were discarded to allow for T1 equilibration effects. Prior to functional scanning, a 6 min T1-weighted MPRAGE structural scan was collected at a resolution of 1×1×1 mm. Stimuli were projected onto a screen behind the participant and viewed in a mirror. Participants responded using a 4-button response box. All stimuli were presented with Cogent running under Matlab2014, permitting synchronisation with the scanner and accurate timing of stimuli presentation.

Data were processed and analysed using SPM12 (www.fil.ion.ucl.ac.uk/spm). The EPI images from all four sessions of each participant were realigned to a mean EPI image for that participant. Images in which the participant moved more than 1.5mm or had rotation of more than 1 degree were visually examined and if seen to contain artefacts were removed from the analysis and replaced with volumes interpolated from the preceding and subsequent images. No participant had artefacts in more than 5% of images. Each participant’s structural image was processed using a unified segmentation procedure combining segmentation, bias correction, and spatial normalization to the MNI template (Ashburner & Friston, 2005). The same normalization parameters were then used to normalize the EPI images. Finally, the images were spatially smoothed to conform to the assumptions of the GLM implemented in SPM12 by applying a Gaussian kernel of 8 mm FWHM.

In order to examine whether the relationship between the participants preferences and those of the confederates was coded in terms of similarity or consistency two general linear models (GLM) were created to enable analysis of the different trial types in our factorial design and to fit the parameters of our RL model. Both GLMs modelled BOLD activation during *outcome* screen for the Sim confederate and Diff confederate separately. Regressors of no interest modelled activity during the *self-choice outcome* screen, the *feedback* phase, the ratings periods, trials where participants failed to make a choice and the residual effects of head motion. In addition, parametric modulators linked to the *outcome* screen regressors allowed us to model the values of our RL parameters on a trial-by-trial basis.

In the Similarity GLM we modelled the similarity prediction error (PE_Sim) and accumulated similarity (AS) between confederate choice and participant choice for each confederate (n=Sim or n=Dif), using the following algorithms:

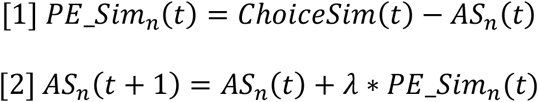

where

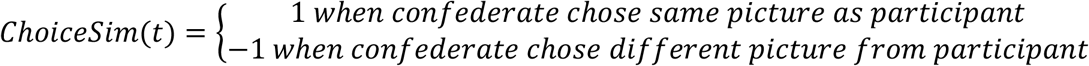

The learning rate (*λ*) was set to 0.5 and initial AS was set to 0 (see Figure 2B for examples of how AS and PE_Sim varied across 20 trials). In total, there were six regressors-of-interest in our Similarity GLM: outcome screens for Sim and Diff; AS values for Sim and Diff, and PE_Sim values for Sim and Diff.

In the Consistency GLM we modelled the prediction error (PE_Con) and accumulated consistency (AC) between confederate choice and participant choice for the two confederates (n=Sim or n=Dif), using the following algorithm.

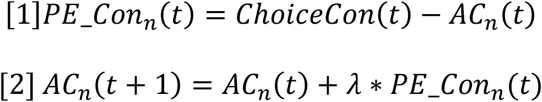

where

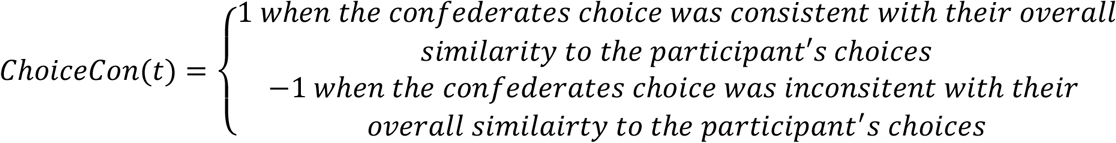

Again, the learning rate (*λ*) was set to 0.5 and initial AC was set to 0 (see Figure 2C for examples of how AS and PE varied across 20 trials). In total, there were six regressors-of-interest in our Consistency GLM: outcome screens for Sim and Diff; AC values for Sim and Diff, and PE_Con values for Sim and Diff.

For each GLM SPM12 was used to compute first level parameter estimates (beta) and contrast images (containing weighted parameter estimates) for each contrast at each voxel. To examine regions showing a main effect of confederate similarity, two contrasts were carried out between the *outcome screen* regressors (Sim > Diff, Diff > Sim). In addition, to examine regions that tracked the RL model parameters in each model, conjunction images were calculated for each RL parameter (AS _Sim_ /AC_Sim_ ∩ AS _Diff_/AC_Diff_) and (PE_Sim _Sim_ /PE_Con _Sim_ ∩ PE_Sim _Diff_/PE_Con _Diff_).

For the group-level analysis, the first level images from all participants were subjected to two one-sample t-tests, one in the positive direction and the other in the negative direction. Images derived from these second level analyses were thresholded at p < 0.001, uncorrected. For each analysis, a separate Monte Carlo simulation implemented in 3dClustSim (Forman et al., 1995) was used to determine the correct cluster extent threshold needed for a whole brain cluster-wise significance level of p < 0.05. Anatomical Regions were determined using the AICHA atlas (Joliot et al., 2015).

## 3 Results

### 3.1 Behavioural Results

To examine whether learning about the preferences of the confederates changed participants feelings of affiliation towards them, we collected ratings of similarity, likeability and trustworthiness at the start of the study and after every 20 trials. These data were then z-scored within participant to normalise them. Separate 2 (confederate: similar/different) × 5 (session number: pre/S1/S2/S3/S4) repeated measures ANOVAs were carried out on the z-scored ratings of similarity, liking and trust (See Figure 3). Due to problems with data recording, the ratings from 7 participants were incomplete and were excluded from the behavioural analysis leaving a remaining sample of 18 participants.

**Figure 3.**
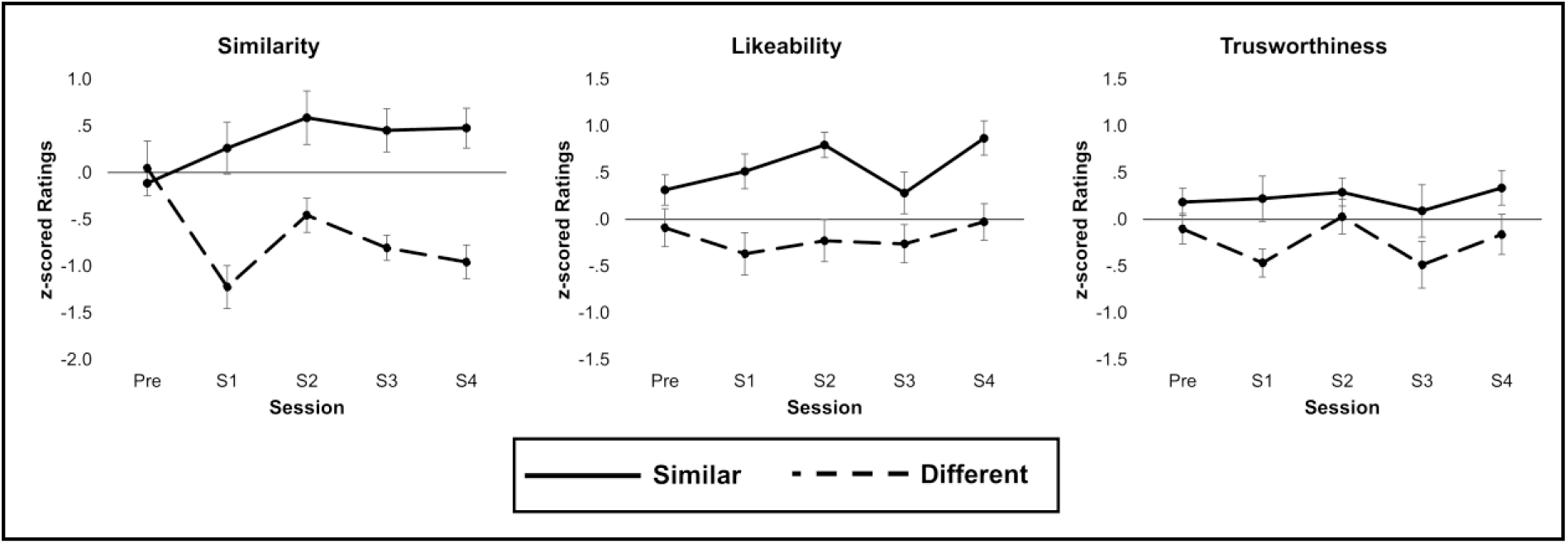
Z-scored ratings of liking similarity and trustworthiness for the similar and different confederates across rating sessions.

The ANOVA on similarity ratings found a significant main effect of confederate, *F*(1,17) = 23.52, *p* < .001, η^2^_p_ = .58. Overall participants rated the similar confederate as being more similar (*M* = 0.33, *MSE* = 0.15) to them than the different confederate (*M* = −0.68, *MSE* = 0.12). There was also a significant interaction between confederate and session *F*(1,17) = 5.65, *p* = .001, η^2^_p_ = .25. To examine this interaction further, ratings for the different confederate was subtracted from the ratings of the similar confederate for each session to create a difference score. Pairwise comparisons (Bonferroni corrected) showed that the difference score for the pre-session (*M* = −0.16, *MSE* = 0.36) significantly differed from the scores after sessions S1 (*M* = 1.49, *MSE* = 0.35), *p* < .05, S3 (*M* = 1.26, *MSE* = 0.29), *p* < .05, and S4 (*M* = 1.43, *MSE* = 0.25), *p* < .01. No other pairwise comparisons were significant.

The ANOVA on liking ratings found a significant main effect of confederate, *F*(1,17) = 23.8, *p* < .001, η^2^_p_ = .58. Overall participants rated the similar confederate as being more likeable (*M* = 0.55, *MSE* = 0.07) than the different confederate (*M* = −0.2, *MSE* = 0.12). There was no significant main effect of session and no interaction between session and confederate. The ANOVA on trust ratings found a significant main effect of confederate, *F*(1,17) = 7.67, *p* < .05, η^2^_p_ = .31. Overall participants rated the similar confederate as being more trustworthy (*M* = 0.23, *MSE* = 0.11 than the different confederate (*M* = −0.24, *MSE* = 0.01). There was no significant main effect of session and no interaction between session and confederate.

### 3.2 fMRI Results

#### 3.2.1 Main Effect of Confederate Preference Similarity

Two key contrasts investigated the effects of confederate identity (Sim/Diff) on BOLD response while participants observed the outcome of confederates’ choices. The regressors which contribute to these contrasts were identical in the Similarity GLM and the Consistency GLM, so the results here are the same for both. The different > similar contrast revealed that observing the choice of the different compared to the similar confederate led to greater activation in the right inferior frontal sulcus (rIFS) and in a cluster centred on the right fusiform gyrus (rFG) (see Table 1 and Figure 4A). No significant activations were found in the similar > different contrast.

**Table 1.** Peak voxel coordinates in MNI space, z-values and cluster sizes for analyses of the choice period showing significant effects after cluster correction for main effect of similarity. Same shading indicates local maxima in distinct anatomical regions within the same cluster, BA indicates Brodman Area, k indicates the cluster size threshold for whole brain significance of *p* < 0.05.

| Region | Hem. | X | Y | Z | Z-Score | Cluster Size |
| --- | --- | --- | --- | --- | --- | --- |
| **Different > Similar (k = 33)** |  |  |  |  |  |  |
| Inferior Frontal Sulcus (BA 44) | R | 38 | 10 | 34 | 3.86 | 59 |
| Fusiform Gyrus (BA 18) | R | 14 | -82 | -10 | 3.51 | 77 |
| Lateral Occipital Gyrus (BA 19) | R | 30 | -82 | -14 | 3.40 |  |

**Figure 4.**
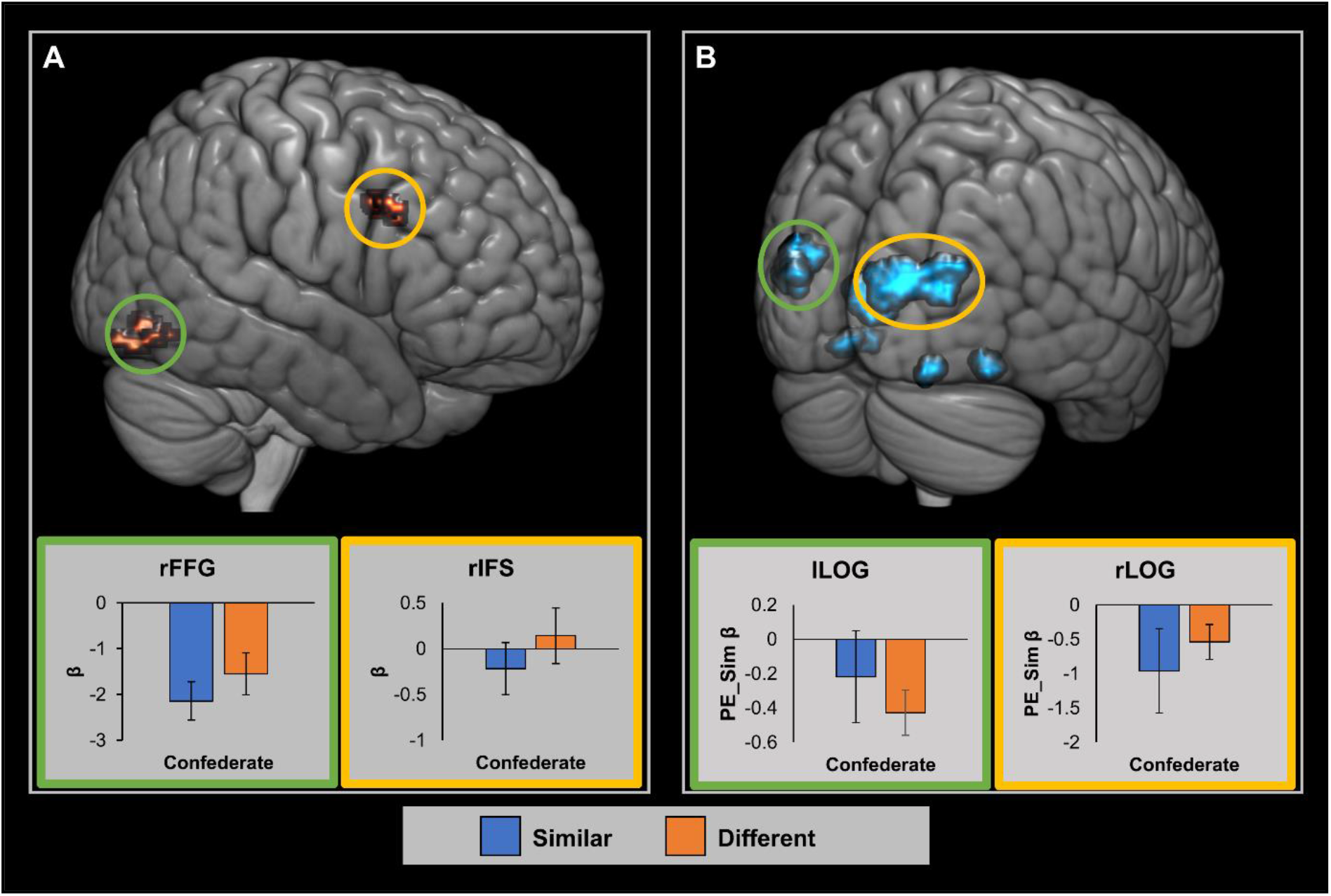
A) Brain areas showing significant cluster corrected results in the Different > Similar contrast. B) Brain areas showing significant cluster corrected results in the negative PE_Sim conjunction. Parameter estimates averaged across whole cluster. Error bars represent SEM. Graph border colours indicate matching circled area. Red/yellow represents positive activations and blue/green represents negative activations.

#### 3.2.2 Parametric Analysis of the Similiarity GLM

To identify brain regions which track acculumated similarity in the Similarity GLM, we calculated a conjunction of the RL parameters for each of the confederates, that is: AS_Sim_ ∩ AS_Diff_. This did not reveal any significant clusters in either a positive or negative direction, suggesting that no brain areas directly tracked preference similarity between confederate and participant. Similarly there were no significant clusters that tracked the positive conjuction of similarity prediction error for both confederates, that is, PE_Sim_Sim_ ∩ PE_Sim_Diff_. This means that no areas showed increased activation when both confederates preference was unexpectedly similar to that of the participant. However, the negative PE_Sim conjunction analysis revealed that unexpected dissimilarity between both confederates and participant choice was positively correlated with activation in a number of clusters within the occipital cortex including the bilateral lateral occipital cortex (LOC) and the lingual gurus (see Table 2 and Figure 4B).

**Table 2.**
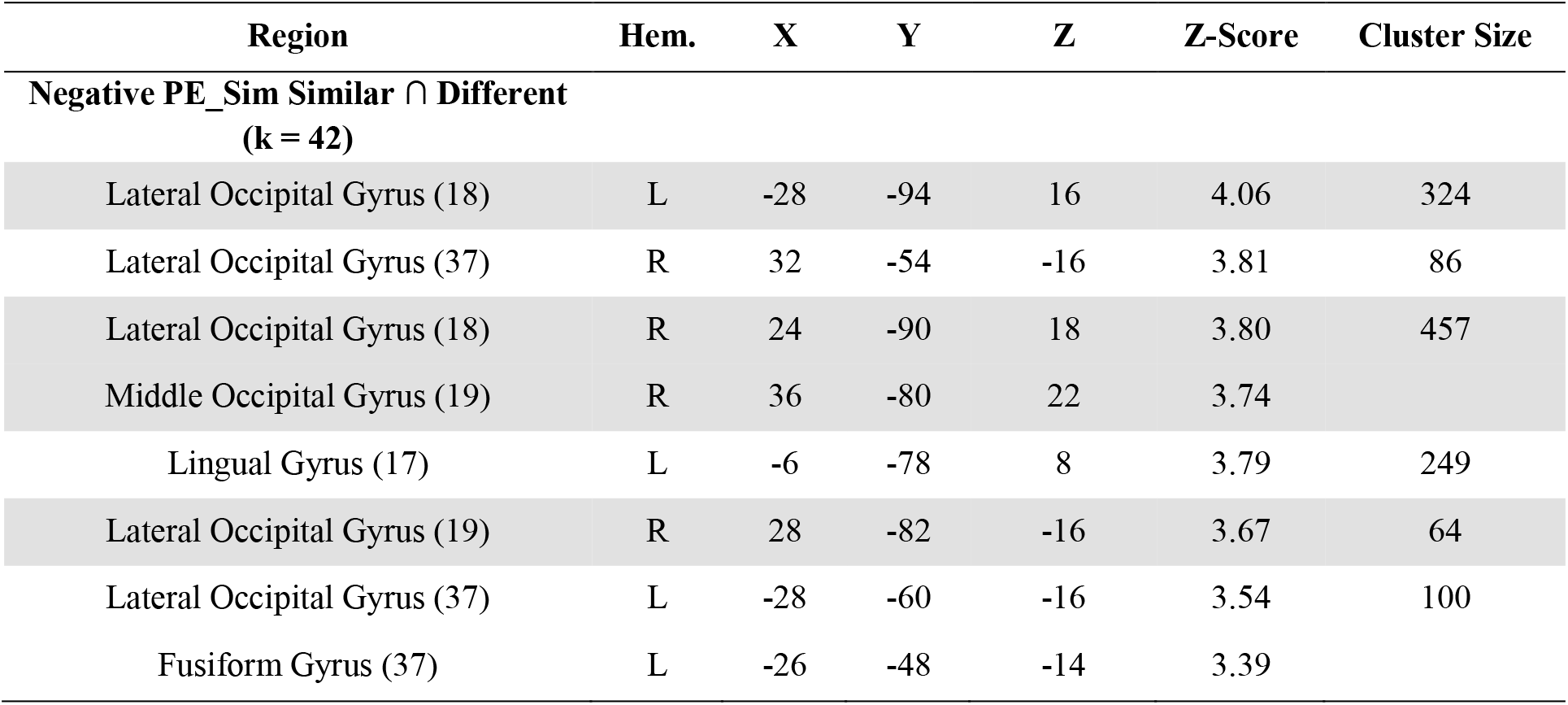
Peak voxel coordinates in MNI space, z-values and cluster sizes for analyses of the choice period in the Similarity GLM showing significant effects after cluster correction for conjunction analyses of the AS and PE parametric modulators. Same shading indicates local maxima in distinct anatomical regions within the same cluster, BA indicates Brodman Area, k indicates the cluster size threshold for whole brain significance of *p* < 0.05.

#### 3.2.3 Parametric Analysis of the Consistency GLM

To identify brain regions tracking the consistency of confederates choices, we first examined the conjunction of areas tracking accumulated consistency for the Similar and Different confederates, that is AC_Sim_ ∩ AC_Diff_. The positive conjunction showed a significant activation in a cluster-corrected region centred on the right superior medial frontal gyrus (smFG) extending into the left superior medial frontal gyrus (see Table 3 and Figure 5A). That means that these areas show greater activation as evidence for the consistency of the confederates’ choices with their previous choices increased, and lower activation during inconsistence periods. No significant activations were found in the conjunction analysis testing for areas negatively correlated with AC.

**Table 3.** Peak voxel coordinates in MNI space, z-values and cluster sizes for analyses of the choice period in the Consistency GLM showing significant effects after cluster correction for conjunction analyses of the AS and PE parametric modulators. Same shading indicates local maxima in distinct anatomical regions within the same cluster, BA indicates Brodman Area, k indicates the cluster size threshold for whole brain significance of *p* < 0.05.

| Region | Hem. | X | Y | Z | Z-Score | Cluster Size |
| --- | --- | --- | --- | --- | --- | --- |
| Positive AC Similar ∩ Different (k = 43) | | | | | | |
| Superior Medial Frontal Gyrus (9) | R | 8 | 56 | 34 | 3.37 | 76 |
| Superior Medial Frontal Gyrus (10) | L | -2 | 54 | 24 | 3.25 |  |
| Superior Medial Frontal Gyrus (10) | R | 6 | 56 | 22 | 3.17 |  |
| Positive PE_Con Similar ∩ Different (k = 43) | | | | | | |
| Caudate Nucleus | L | -12 | -6 | 28 | 4.54 | 52 |
| Caudate Nucleus | R | 16 | -6 | 28 | 3.90 | 71 |
| Caudate Nucleus | L | -4 | 14 | 12 | 3.64 | 56 |
| Negative PE_Con Similar ∩ Different (k = 43) | | | | | | |
| Angular Gyrus (40) | R | 56 | -46 | 50 | 4.22 | 341 |
| Interparietal Sulcus (40) | R | 32 | -50 | 40 | 3.70 |  |
| Superior Frontal Sulcus (10) | R | 34 | 50 | 10 | 4.20 | 270 |
| Superior Temporal Sulcus (37) | R | 60 | -58 | 16 | 3.85 | 76 |
| Superior Temporal Sulcus (41) | R | 44 | -42 | 20 | 3.72 | 43 |
| Superior Temporal Sulcus (39) | R | 42 | -54 | 16 | 3.19 |  |
| Precuneus (39) | R | 10 | -56 | 48 | 3.72 | 106 |
| Middle Temporal Gyrus (21) | R | 60 | -20 | -16 | 3.46 | 57 |
| Superior Temporal Sulcus (21) | R | 62 | -28 | -10 | 3.43 |  |

**Figure 5.**
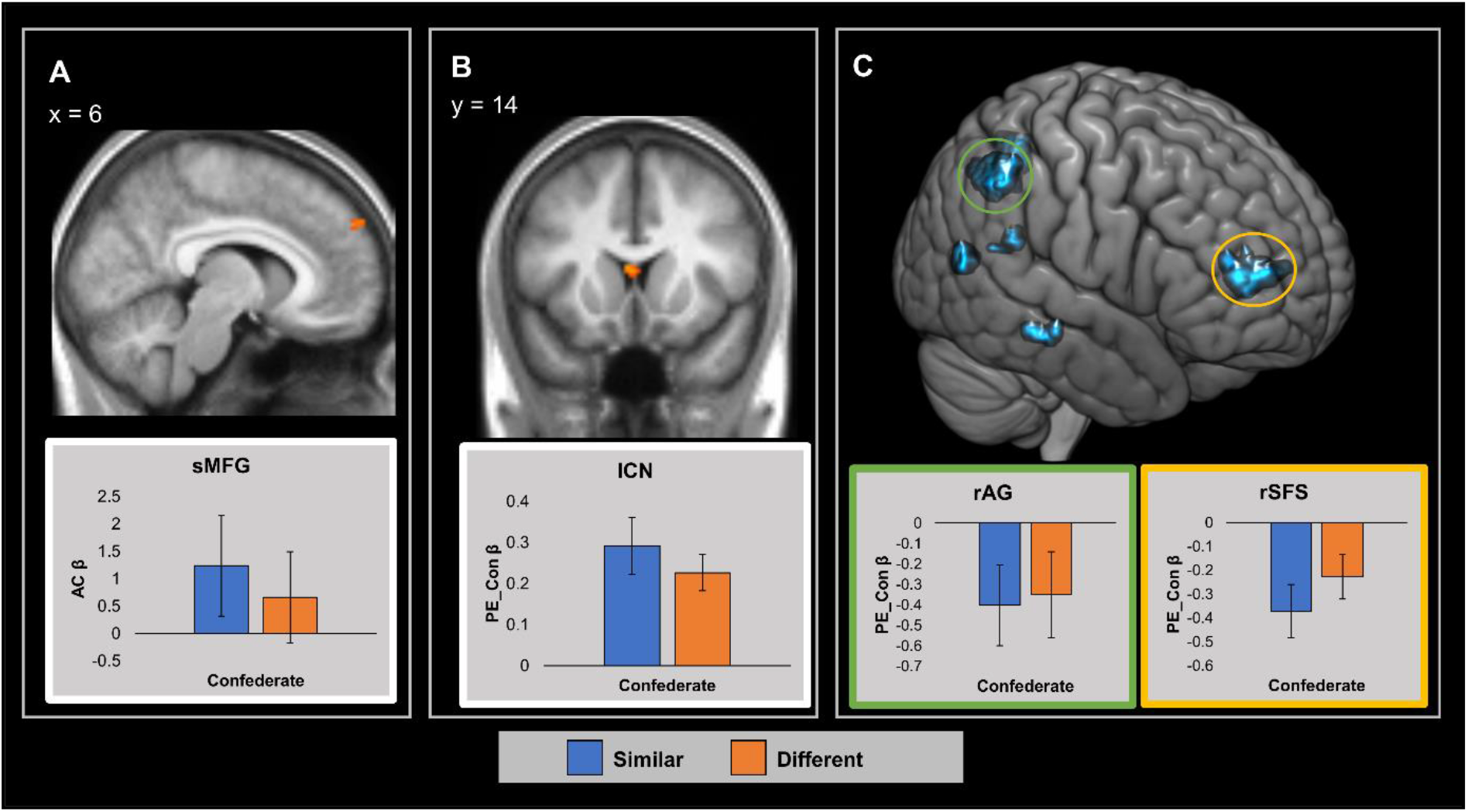
Brain areas showing significant cluster corrected tracking of AC and PE_Con. A) Areas significantly tracking AC in the positive CSim ∩ CDiff conjunction. B) Areas significantly tracking PE_Con in the positive CSim ∩ CDiff conjunction. C) Areas significantly tracking PE_Con in the negative CSim ∩ CDiff conjunction. Parameter estimates averaged across whole cluster. Error bars represent SEM. Graph border colours indicate matching circled area. Red/yellow represents positive activations and blue/green represents negative activations. sMFG = superior Medial Frontal Gyrus, lCN = left Caudate Nucleus, rAG = right Angular Gyrus, rSFS = right Superior Frontal Sulcus.

The conjunction analysis testing for areas tracking prediction error in consistency (PE_Con_Sim_ ∩ PE_Con_Diff_) identified significant cluster-corrected activations bilaterally in a dorsal region of the caudate nucleus as well as in a more ventral region of the left caudate nucleus (see Table 3 and Figure 5B). That means that these areas showed increased BOLD response when the confederates’ choice was unexpectedly consistent with their overall preference, and decreased activation when confederates’ choice was unexpectedly inconsistent. Note that while the peak activations in the two dorsal clusters are in fact found in the neighbouring ventricles all three clusters show considerable overlap with the caudate nucleus. The conjunction analysis testing for areas tracking PE_Con in a negative direction identified significant clusters in several right hemisphere regions, namely the angular gyrus (rAG), the superior frontal sulcus (rSFS), the superior temporal sulcus (rSTS), the medial temporal gyrus (rMTG) and the Precuneus (see Table 3 and Figure 5C). That means these areas show increased BOLD response when the confederates’ choice was unexpectedly inconsistent with their overall preference, and reduced activity when the confederates’ choice was highly predictable.

## 4 Discussion

Our study examined the neural basis of learning about preference similarity between self and other and its role in promoting affiliation. We created a context where participants could express a preference for a painting and learn about the preferences of two confederates for the same paintings. Our behavioural data supports the claim that similar preferences lead to higher ratings of liking, trustworthiness and similarity. This behavioural result indicates that participants were affected by our experimental manipulation, and therefore tracked the confederates’ preferences in relation to their own preferences. To track the brain patterns underlying this preferences we modelled our neuroimaging data with a Similarity model which tracks how similar the confederates are to the participant, and a Consistency model which tracks how consistent each confederate is with their previous behaviour in a person-specific fashion. We found several regions linked to reward and social cognition which track parameters of the Consistency model. This finding suggests that trial-by-trial social learning about others’ preference similarity is largely implemented on a person-specific basis and not on their similarity to the self.

Our introduction outlined two ways that preference similarity might be tracked for similar and different others: either by a general mechanism which tracks all people in relation to how close or distant they are to oneself, or via a person-specific mechanism which tracks similarity in terms of how consistent a person’s current choice similarity is to their previous similarity to the self. To examine the evidence for these two mechanisms we created two RL models which tracked confederate’s choices based on similarity and consistency respectively. Our results from the similarity model indicated that unexpectedly dissimilar choices from either participant correlated with increase activation in several regions of the visual cortex. Results from the consistency model showed a greater variety of regions with a region of the dmPFC tracking AC for both participants while activity in a range of regions involved in domain general reward learning and social cognition tracked consistency prediction errors. Below we elaborate on the results of the AC conjunction before moving on to discuss the findings on PE_Con and PE_Sim.

### 4.1 dmPFC Implicated in Representing the Consistency of Preference Similarity between Self and Other

The accumulated consistency (AC) parameter represents a trial-by-trial estimate of the probability that a confederate makes choices similar to their previous choices, that is, that the Sim confederate should choose the same painting as the participant while the Diff confederate should choose differently. The only area we found tracking AC was a cluster in the bilateral smFG corresponding to the anterior region of the dmPFC. The dmPFC is known to be a key area for the processing of information about both self and other (Amodio & Frith, 2006; Eickhoff, Laird, Fox, Bzdok, & Hensel, 2014; Mitchell, Banaji, & Macrae, 2005). To place our findings in context, we identified 15 studies that had found activation in the dMPFC and categorised the contrasts those activations came from into the following four categories: 1) Diagnostic > Non-Diagnostic cases: contrasting information relevant to a trait judgement about an individual with irrelevant information; 2) Inconsistent > Consistent: cases contrasting novel information that was inconsistent with previous knowledge about an individual with novel information that was consistent with previous knowledge; 3) Other Impression Formation: other contrasts relevant to impression formation, often linking photographs of individuals with information about their traits; 4) Self Relevant: contrasts in which participants judged whether information was self-relevant or not (see Supplementary Material for further details). We then collapsed the clusters found in these studies across the x-axis to create Figure 4. Our result falls in the middle of the region activated by previous studies, with most contrasts investigating Inconsistent > Consistent falling more dorsally and most contrasts investigating Diagnostic > Non-Diagnostic falling more ventrally suggesting that the activation found in our study is in agreement with the previous literature.

The involvement of the dmPFC in coding prior knowledge of other is supported by previous research suggesting that the dmPFC encodes reputational priors of one’s partners during economic games (Fouragnan et al., 2013; Hampton, Bossaerts, & O’Doherty, 2008). Our results build on these findings by suggesting that the dmPFC represents the strength of prior evidence in the context each confederate’s overall pattern of preference similarity rather than simply displaying increased activation for similar preferences regardless of confederate identity.

**Figure 6.**
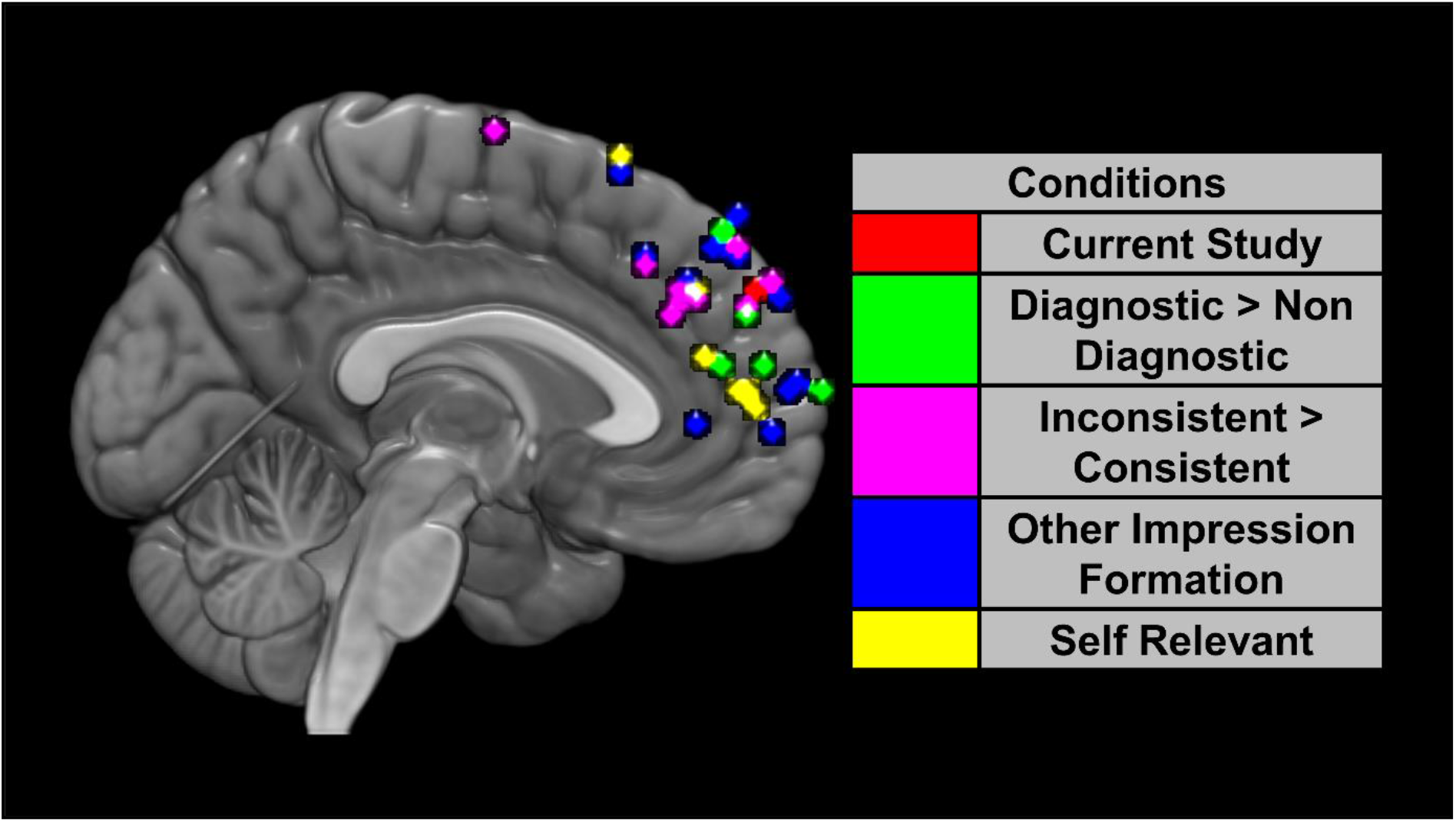
dmPFC activations across 15 studies investigating impression formation and the self together with the result from the current study. For ease of presentation we have collapsed the studies across the x-axis (max x = 14, min x = −13).

### 4.2 Consistency Prediction Errors Tracked by Regions Involved in Reward and Social Cognition

The consistency prediction error parameter (PE_Con) reflects the difference between the choice a confederate makes on a specific trial and the participant’s expectation of what choice the confederate will make, based on the confederate’s past behaviour. For example, the model assigns positive update signal when the different confederate picks the painting not chosen by the participant, and a high and negative prediction error is seen when the different confederate picks the same painting. For the similar confederate the same model assigns negative prediction error when the confederate picks the painting not chosen by the participant, and a positive signal when the confederate picks the same painting (See Figure 2). Examination of areas that tracked PE_Con indicated two distinct patterns of activations across different regions. Clusters in the bilateral caudate nucleus (Figure 4B) showed increased activity when confederates chose consistent with their type and decreased acrtivity when they chose against type. Clusters in regions associated with social cognition including the STS, SMG, Precuneus and SFS (Figure 4C) showed increased activations when the confederates choice was inconsistent with their overall pattern of similarity to the participant. Overall, this pattern shows that PE tracking in these regions is not a ‘generic’ signal of how similar a person is to me, but rather tracks specifically how each person’s choice conforms to or deviates from their typical pattern of similarity to me. We now consider how these findings relate to previous studies of these areas.

The caudate nucleus, along with other parts of the striatum, has been heavily implicated in the generation of prediction errors during reinforcement learning of rewards for self (Balleine, Delgado, & Hikosaka, 2007; O’Doherty et al., 2004; Schultz, 2015) and others (Báez-Mendoza & Schultz, 2013; Bhanji & Delgado, 2014; Ruff & Fehr, 2014). Previous studies have shown that the caudate nucleus is also involved in signalling prediction errors when learning the characteristics of others. King-Casas et al. (2005) found that the caudate nucleus activity tracked PEs regarding the trustworthiness of other during an economic game. Subsequent studies have found similar results for trustworthiness (Fareri, Chang, & Delgado, 2012; Fett, Gromann, Giampietro, Shergill, & Krabbendam, 2014; Fouragnan et al., 2013) and that the caudate nucleus also produces PEs when learning about other’s generosity (Fareri et al., 2012), reliability in advice giving (Diaconescu et al., 2017) and general behavioural traits (Mende-Siedlecki & Todorov, 2016). Our findings add to this literature by showing that the caudate nucleus is also involved in learning about the similarity of one’s preferences to the preferences of others.

The regions showing greater activations when PE_Con was negative, i.e. when the confederate choice was inconsistent with their overall pattern of choice similarity, are key nodes of the mentalising network involved in processing information about self and others (Barrett & Satpute, 2013; Murray, Debbané, Fox, Bzdok, & Eickhoff, 2015; Spreng, Mar, & Kim, 2009; Van Overwalle, 2009). These areas have been implicated in the formation of impressions about other peoples’ traits (Gilron & Gutchess, 2012; Hackel et al., 2015; Hughes et al., 2017; Ma et al., 2012; Mende-Siedlecki, Cai, et al., 2013), beliefs (Cloutier et al., 2011) and abilities (Bhanji & Beer, 2013; Mende-Siedlecki, Baron, et al., 2013). Of particular note are two studies which directly modelled PEs for learning about the traits of other. Hackel et al. (2015) found that the precuneus and STS tracked PEs for confederate generosity during an economic game while Stanley (2016) found that only the precuneus showed greater tracking of PEs in a social verses non-social setting. The current study expands on these previous findings by showing that these regions also track PEs regarding the similarity relationship between self and others thus underlining the important role of prediction error in social learning (Joiner et al., 2017).

It is also notable that while previous studies on social impression formation have tended to show bilateral activations of the mentalising network the current studies activity was limited to the right hemisphere. This is consistent with previous research demonstrating right lateralisation for tasks involving self and other differentiation (Decety, 2003; Hu et al., 2016; Kaplan, Aziz-Zadeh, Uddin, & Iacoboni, 2008; Uddin, Kaplan, Molnar-Szakacs, Zaidel, & Iacoboni, 2005).

### 4.4 Similarity Related Responses in Regions Involved in Visual Attention

In addition to modelling the RL parameters, we also directly contrasted the outcome screen where participants see the choices of the similar confederate with the outcome screen for the different confederate. This contrast shows greater activation for the different confederate in two clusters; one centred on the rIFS and the other on the rFG. The IFS has been implicated in attentional processing and in particular in the control of attentional shifts by both internal goals and by salient external stimuli (Aron, Robbins, & Poldrack, 2004, 2014; Asplund, Todd, Snyder, & Marois, 2010; Filimon, Philiastides, Nelson, Kloosterman, & Heekeren, 2013; Levy & Wagner, 2012), while the FG is known to play a key role in the visual perception of faces (Contreras, Banaji, & Mitchell, 2013; Kanwisher & Yovel, 2006; Rotshtein, Henson, Treves, Driver, & Dolan, 2005). In the current study, the faces of the confederates were displayed on the screen during the outcome period. Interestingly a previous study also found greater FG activation when participant observed faces of individuals judged to have different traits to themselves (Leshikar, Cassidy, & Gutchess, 2016).

The activation of these areas suggests that participants may have found the choices of different confederate to be more attention grabbing than those of the similar confederate in a comparable way to studies that have demonstrated an attentional bias towards untrustworthy as opposed to trustworthy others (Dzhelyova, Perrett, & Jentzsch, 2012; Farmer, Apps, & Tsakiris, 2016; Vanneste, Verplaetse, Van Hiel, & Braeckman, 2007). These findings were also consistent with our conjunction analysis of regions that showed a negative relationship to the value of PE_Sim. This analysis revealed that when a confederate made an unexpectedly dissimilar choice to that of the participant it led to increase activation across a series of visual areas including regions in the bilateral LOC and in the left FFG. Again these results support the idea that different choices may be more visually attention grabbing than similar ones leading to greater activation in visual cortices.

### 4.5 Limitations and Future Directions

One key limitation of the current study is that our task did not allow us to collect trial-by-trial behavioural data showing what participants had learnt about the confederates. This is because we wanted participants to learn implicitly, rather than making explicit predictions of the confederate’s choice on each trial. As we did not collet trial-by-trial responses data, we approximated a preference learning rate (0.5) and used it in our RL models to track changes in preference tracking according to the actual choices made by the confederates. This raises the possibility that there may only be a weak fit between the learning rate used in our model and the actual learning rate of our participants. However, our main predictions and results had to do with the direction of the tracked prediction errors and accumulated preferences, and not with the specific magnitude of these variables, which are less likely to be affected by our approximation. This is in line with a recent theoretical paper (Wilson & Niv, 2015) that demonstrated that model based fMRI results are, under some conditions, insensitive to changes in individual learning rates While it is possible that our approximation may be lead to lower power at detecting brain responses to prediction errors, we feel that the main hypothesis concerning the direction of the effects (Similarity approach vs Consistency approach) is supported by our analysis. However, the lack of behavioural data does constrain our ability to test different models of preference learning in a more structured manner. Future studies could expand upon our work by asking participants to explicitly predict the choices of the confederates which would allow for more detailed testing of different computational models but might also alter the social nature of the task.

A second limitation of the current study relates to the motivation underlying our participants’ behaviour. Our participants received no reward based on their choices which may have limited the sense that the choices made were meaningful. It is noticeable that previous studies that gave monetary rewards (Garvert et al., 2015) or rewards in kind (Campbell-Meiklejohn et al., 2010) have shown VS activation when tracking similarity between their own choices and those of others. By contrast no VS activation were seen in the current study nor in previous preference comparison studies in which the participant did not receive a meaningful reward for either their own choice or those of others (Welborn et al., 2016). However, we note that our study (without monetary rewards) gives a better match to real-world contexts where the choices that we and other people make in our social interactions often do not lead to immediate financial benefits.

One final limitation of the current study is that, due to the use of a multiband imaging sequence, it was not possible to analyse the connectivity between brain areas, for example those identified as tracking PE_Con and the dmPFC. In our case multiband was necessary for us to achieve a fine enough TR so separately model the learning of both the similar and different confederates but with a different design it would be possible to create a similar setup in which learning for similar and different others took place in separate trials allowing for a more traditional scanning protocol that is amenable to functional connectivity modelling.

Further research in this area could build on our results by examining whether the neural correlates of similarity learning are modulated by having pre-existing cues about how similar that person is to oneself. For example, does learning that someone has similar or different values to oneself modulate how quickly one learns about their preferences in another domain such as art or music and is prior similarity represented in the same way across those domains? The relationship between preference similarity and social influence would be another interesting direction for further research. The areas shown to be activated by similarity learning in the current study strongly overlap with those implicated in preference shifting based on social influence in previous studies (Campbell-Meiklejohn et al., 2010; Welborn et al., 2016). It would be interesting to see whether participants were more likely to shift their preferences in the direction of another person once they have learnt that person generally shares their preferences and if this process is mediated by the neural mechanisms involved in similarity learning.

Finally, our current study found evidence that we track other people in terms of their consistency with their previous behaviour, which means that each confederate is tracked as a distinct individual with particular preferences. In the present project, consistency was defined in relation to self-choices, that is, the Sim confederate tended to choose the same as the participant and the Diff confederate tended to make a different choice. We think it likely that a consistency model would also apply if confederates made consistent choices on a different dimension, for example, if A consistently chose blue pictures and B consistently chose red pictures. However, further work would be needed to test this and reveal how much this consistency approach applies to learning about other domains including people’s traits, attitudes and competence.

### 4.6 Conclusions

In this study, we combined computational modelling and fMRI to investigate the neural processes that underlie learning about the similarity of other’s preferences to one’s own. We found that the brain encodes the similarity of others’ choices in a consistency driven manner with brain areas that represent both accumulated preference consistency and consistency prediction errors, and not in a manner dependent on the others’ similarity to the oneself. These findings indicate that the neural representations of similarity to the self are coded in a person specific manner that reflects how consistent and inconsistent the similarity that person’s current preference similarity to the self is with the overall similarity of their past preferences. By doing so they highlight the role of context dependent predictive processing in the learning of preference similarity between self and other and, by extension, in the formation of social impressions more generally.

## Funding

This work was supported by European Research Council (http://erc.europa.eu/) grant 313398-INTERACT.

## Acknowledgements

The authors acknowledge the help of Ms Juliette Klamm in the validation of the experimental stimuli and pilot testing for the study and Ms Xijing Wang in the collection of fMRI data for the study.

