## Supplementary Materials for "The Neural Basis of Shared Preference Learning"

### The Neural Basis of Shared Preference Learning – Supplementary Materials

#### S1. Supplementary Methods

##### S1.1. Materials

The picture stimuli used in this study comprised of 40 pairs of abstract paintings and 40 pairs of landscape paintings that were matched as closely as possible in terms of their visual and aesthetic properties. This was done to ensure that different pairs of paintings would not differ wildly in their characteristics so that the choices of the different confederate would not suggest an abnormal set of preferences. To construct these sets 120 abstract and 120 landscape images were downloaded from the internet and resized to 390x390 JPEG images with any remaining space on either dimension filled in with black. These images were then rated in a pre-study by a group of 20 participants on their complexity, concreteness, attractiveness, valence, affectivity and interest using a 7-point scale. In addition each images' luminance and contrast were calculated using MATLAB (Mathworks 2015).

The mean ratings and luminance and contrast measures for each image were standardised across all images. Similarity scores were created for each measure by subtracting each images score from the score of every other image of its group (abstract or landscape). The similarity scores for all measures were then combined into one value using the following algorithm:

$$(Attractiveness * 2) + \ (Interest * 2) + Complexity + Valance + Affectivity + \left(\frac{Concreteness}{2}\right) + \left(\frac{Luminance}{2}\right) + \left(\frac{Contast}{2}\right)$$

Each of the images was then paired with its closest neighbour and each pair was then removed from the array. The 40 closest pairs in each group were used in the fMRI experiment and the next 5 closest pairs were used in the training block.

#### S2. Details of dmPFC literature comparison

**Table S2.** Details of studies used in the comparison of dmPFC activation across impression formation studies (see Figure 8). Coordinates are reported in MNI space and coordinates from studies using Talairach space were converted using the WFU PickAtlas version 2.4 (Maldjian, Laurienti, and Burdette 2004; Maldjian et al. 2003).

| Study | Information presented | Impression studied | X | Y | Z |
|---|---|---|---|---|---|
| **Current Study (Red)** | Choice of preferred painting | Similarity to Self | 8 | 5 | 34 |
| **Diagnostic > Non Diagnostic (Green)** | | | | | |
| Gilron & Gutchess, 2012 | Moral and Neutral Behaviours | Moral Impression vs Location of Behaviour | 0 | 57 | 15 |
| | Moral and Neutral Behaviours | Moral Impression vs Location of Behaviour | 3 | 48 | 48 |
| | Moral and Neutral Behaviours | Moral Impression vs Location of Behaviour | 9 | 72 | 9 |
| Ma, Vandekerckhove, Van Overwalle, Seurinck, & Fias, 2010 | Moral and Neutral Behaviours | Personality Traits | 4 | 54 | 28 |
| | Moral and Neutral Behaviours | Personality Traits | 14 | 48 | 16 |
| **Inconsistent > Consistent (Pink)** | | | | | |
| Cloutier, Gabrieli, Young, & Ambady, 2011 | Political Views | Political Affiliation | 2 | 54 | 30 |
| Cloutier et al., 2011 | Political Views | Political Affiliation | 6 | 52 | 44 |
| Ma et al., 2012 | Moral vs Neutral Behaviours | Moral Traits | 4 | 42 | 32 |
| | Moral vs Neutral Behaviours | Moral Traits | 4 | 38 | 34 |
| | Moral vs Neutral Behaviours | Moral Traits | 4 | 35 | 28 |
| Mende-Siedlecki, Cai, & Todorov, 2013 | Moral Behaviours | Moral Impession | 2 | 31 | 40 |

| | | | | | |
|---|---|---|---|---|---|
| Mende-Siedlecki & Todorov, 2016 | Moral and Neutral Behaviours | Trustworthiness and Surprise | -5 | 59 | 36 |
| | Moral and Neutral Behaviours | Trustworthiness and Surprise | -2 | -5 | 72 |
| **Other Impression Formation (Blue)** | | | | | |
| Ames & Fiske, 2013 | Assessment of Teaching Ability | Expertise | -4 | 45 | 45 |
| Baron, Gobbini, Engell, & Todorov, 2011 | Moral Behaviours | Trustworthiness | -4 | 41 | 36 |
| Freeman, Schiller, Rule, & Ambady, 2010 | Individuated vs Superficial for Racial In-group vs Out-group | Personality Traits | -13 | 43 | 4 |
| Fouragnan et al., 2013 | Choices in Trust Games and Prior information about Trustworthiness | Trustworthiness | -2 | 64 | 10 |
| | Choices in Trust Games and Prior information about Trustworthiness | Trustworthiness | 0 | 62 | 31 |
| Gilron & Gutchess, 2012 | Moral Behaviours vs Neutral Behaviours | Moral Impression vs Location of Behaviour | 3 | 30 | 42 |
| Mende-Siedlecki, Baron, & Todorov, 2013 | Moral and Ability Behaviours | Competence and Trusworthiness | -5 | 66 | 13 |
| | Moral and Ability Behaviours | Competence and Trusworthiness | 32 | 60 | -1 |
| Mende-Siedlecki, Cai, et al., 2013 | Moral Behaviours | Moral Impression | 5 | 52 | 51 |
| Schiller, Freeman, Mitchell, Uleman, & Phelps, 2009 | Moral Behaviours | Moral Impression | -9 | 24 | 61 |
| Schiller et al., 2009 | Moral Behaviours | Moral Impression | -7 | 52 | 43 |
| **Self (Yellow)** | | | | | |
| Martinelli et al. 2013 | Memory Meta-Analysis | Semantic Autobiographic Memory | -10 | 45 | 18 |
| | Memory Meta-Analysis | Episodic Autobiographic Memory | -6 | 51 | 9 |
| | Memory Meta-Analysis | Conceptual Self | 6 | 55 | 5 |
| Moran, J.M. et al., 2006 | Personality Traits | Self-Relevance | -6 | 53 | 6 |

| | | | | | |
|---|---|---|---|---|---|
| Phan, K.L. et al., 2004 | Emotional Pictures | Self-Relatedness | 0 | 42 | 33 |
| Schneider et al. 2008 | Emotional or Neutral Pictures | Self-Relatedness | -3 | 24 | 66 |

Brett, Matthew, Jean-Luc L Anton, Romain Valabregue, and Jean-Baptiste Poline. 2002. "Region of Interest Analysis Using an SPM Toolbox." *NeuroImage* 16 (2.1): 1140.

Cloutier, Jasmin, J D E Gabrieli, D O Young, and Nalini Ambady. 2011. "An fMRI Study of Violations of Social Expectations: When People Are Not Who We Expect Them to Be." *NeuroImage* 57 (2): 583–88.

Fouragnan, Elsa, Gabriele Chierchia, Susanne Greiner, Remi Neveu, Paolo Avesani, and Giorgio Coricelli. 2013. "Reputational Priors Magnify Striatal Responses to Violations of Trust." *Journal of Neuroscience* 33 (8): 3602–11.

Freeman, Jonathan B, Daniela Schiller, Nicholas O Rule, and Nalini Ambady. 2010. "The Neural Origins of Superficial and Individuated Judgments about Ingroup and Outgroup Members." *Human Brain Mapping* 31 (1): 150–59.

Gilron, Roee, and Angela H Gutchess. 2012. "Remembering First Impressions: Effects of Intentionality and Diagnosticity on Subsequent Memory." *Cognitive, Affective and Behavioral Neuroscience* 12 (1): 85–98.

Ma, Ning, Marie Vandekerckhove, Kris Baetens, Frank Van Overwalle, Ruth Seurinck, and Wim Fias. 2012. "Inconsistencies in Spontaneous and Intentional Trait Inferences." *Social Cognitive and Affective Neuroscience* 7 (8): 937–50.

Ma, Ning, Marie Vandekerckhove, Frank Van Overwalle, Ruth Seurinck, and Wim Fias. 2010. "Spontaneous and Intentional Trait Inferences Recruit a Common Mentalizing Network to a Different Degree: Spontaneous Inferences Activate Only Its Core Areas." *Social Neuroscience* 6 (2): 123–38.

Maldjian, Joseph A, Paul J Laurienti, and Jonathan H Burdette. 2004. "Precentral Gyrus Discrepancy in Electronic Versions of the Talairach Atlas." *NeuroImage* 21 (1): 450–55.

Maldjian, Joseph A, Paul J Laurienti, Robert A Kraft, and Jonathan H Burdette. 2003. "An Automated Method for Neuroanatomic and Cytoarchitectonic Atlas-Based Interrogation of fMRI Data Sets." *NeuroImage* 19 (3): 1233–39.

Martinelli, Pénélope, Marco Sperduti, and Pascale Piolino. 2013. "Neural Substrates of the Self-Memory System: New Insights from a Meta-Analysis." *Human Brain Mapping* 34 (7): 1515–29.

Mathworks. 2015. "Matlab R2015b." Natick, MA: The Mathworks Inc.

Mende-Siedlecki, Peter, Sean G Baron, and Alexander Todorov. 2013. "Diagnostic Value Underlies Asymmetric Updating of Impressions in the Morality and Ability Domains." *Journal of Neuroscience* 33 (50): 19406–15.

Mende-Siedlecki, Peter, Yang Cai, and Alexander Todorov. 2013. "The Neural Dynamics of Updating Person Impressions." *Social Cognitive and Affective Neuroscience* 8 (6): 623–31.

Mende-Siedlecki, Peter, and Alexander Todorov. 2016. "Neural Dissociations between Meaningful and Mere Inconsistency in Impression Updating." *Social Cognitive and Affective Neuroscience* 11 (9): 1489–1500.

Moran, Joseph M, C Neil Macrae, Todd F Heatherton, C L Wyland, and W M Kelley. 2006. "Neuroanatomical Evidence for Distinct Cognitive and Affective Components of Self." *Journal of Cognitive Neuroscience* 18 (9): 1586–94.

Phan, K Luan, Stephan F. Taylor, Robert C. Welsh, Shao Hsuan Ho, Jennifer C. Britton, and Israel Liberzon. 2004. "Neural Correlates of Individual Ratings of Emotional Salience: A Trial-Related fMRI Study." *NeuroImage* 21 (2): 768–80.

Schiller, Daniela, Jonathan B Freeman, Jason P Mitchell, James S Uleman, and Elizabeth A

Phelps. 2009. “A Neural Mechanism of First Impressions.” *Nature Neuroscience* 12 (4): 508–14.

Schneider, Felix, F. Bermpohl, A. Heinzel, M. Rotte, M. Walter, C. Tempelmann, C. Wiebking, H. Dobrowolny, H. J. Heinze, and G. Northoff. 2008. “The Resting Brain and Our Self: Self-Relatedness Modulates Resting State Neural Activity in Cortical Midline Structures.” *Neuroscience* 157 (1): 120–31.
